# Conversion of a soluble protein into a potent chaperone *in vivo*

**DOI:** 10.1101/284802

**Authors:** Soon Bin Kwon, Kisun Ryu, Ahyun Son, Hotcherl Jeong, Keo-Heun Lim, Kyun-Hwan Kim, Baik L. Seong, Seong Il Choi

**Affiliations:** Department of Biotechnology, College of Life Science and Biotechnology, Yonsei University, Seoul 03722, Republic of Korea; Department of Pharmacy, Ewha Womans University, Seoul 03760, Republic of Korea; Department of Pharmacology, Center for Cancer Research and Diagnostic Medicine, IBST, School of Medicine, Konkuk University, Seoul 05029, Republic of Korea; Vaccine Translational Research Center (VTRC), Yonsei University, Seoul 03722, Republic of Korea; Department of Biochemistry and Biophysics, Stockholm University, SE-106 91 Stockholm, Sweden.

## Abstract

Protein-folding assistance and aggregation inhibition by cellular factors are largely understood in the context of molecular chaperones. As an alternative and complementary model, we previously proposed that, in general, soluble cellular macromolecules including chaperones with large excluded volume and surface charges exhibit the intrinsic chaperone activity to prevent aggregation of their connected polypeptides, irrespective of the connection types, and thus to aid productive protein folding. As a proof of concept, we here demonstrated that a model soluble protein with an inactive protease domain robustly exerted chaperone activity toward various proteins harboring a short protease-recognition tag of 7 residues in *Escherichia coli*. The chaperone activity of this protein was similar or even superior to that of representative *E. coli* chaperones *in vivo*. Furthermore, *in vitro* refolding experiments confirmed the *in vivo* results. Our findings revealed that a soluble protein exhibits the intrinsic chaperone activity, which is manifested, upon binding to aggregation-prone proteins. This study gives new insights into the ubiquitous chaperoning role of cellular macromolecules in protein-folding assistance and aggregation inhibition underlying the maintenance of protein solubility and proteostasis *in vivo*.

## Introduction

How metastable proteins with respect to aggregation fold efficiently and maintain their solubility in the crowded cellular environment has been a fundamental yet unsolved question in biology [1–4]. Protein aggregation is closely associated with the numerous human disorders, including neurodegeneration [5]. Molecular chaperones and principles underlying their action mechanisms have provided the conceptual frameworks for understanding of protein-folding assistance, aggregation inhibition, and proteostasis *in vivo* [6, 7]. However, chaperones are ineffective for numerous aggregation-prone proteins [8], and they need to be understood with caveats, given accumulating evidence. Chaperones generally assist protein folding by preventing off-pathway “intermolecular” aggregation (or increasing the final folding yield), often at the expense of the “intramolecular” folding rate or thermodynamic stability of substrate proteins [9–11]. By contrast, in some cases, chaperones can increase the folding rate of client proteins as folding catalysts [12–14]. Both action mechanisms of chaperones are basically different although they are not mutually exclusive.

Chaperones commonly recognize and bind to the exposed hydrophobic residues of non-native or unfolded polypeptides, thereby enabling protein quality control [3, 6, 7]. These findings led to widespread beliefs in 1) the hydrophobic interaction-mediated substrate recognition of chaperones; and 2) such interaction-mediated substrate stabilization against aggregation [3, 4, 7, 15]. However, several lines of evidence challenge these prevailing mechanisms. Chaperones, such as Spy and Trigger Factor (TF), can recognize and bind to the surface-exposed charged regions of substrates [16, 17], and the endoplasmic reticulum lectin chaperones calnexin/calreticulin bind to the carbohydrate parts of their substrate proteins [18]. Moreover, GroEL and TRiC/CCT can also recognize their substrates mainly through electrostatic interactions [19, 20]. These findings naturally raise a fundamental question regarding how chaperones stabilize their substrates, which are connected via non-hydrophobic interactions, against aggregation. Contrary to such widespread belief, it remains poorly defined what forces (or factors) of chaperones or other cellular macromolecules are responsible for stabilizing their bound substrates against aggregation. This is primarily due to the inherent difficulty of this study, including conformational changes of macromolecules and irreversible aggregation. Intriguingly, the surface-charge patches of heat-shock protein 90 (HSP90) are critical for the anti-aggregation activity for its substrate proteins, although the charge patches are located away from the substrate-binding regions [21]. Similarly, the substrate-stabilizing ability of HSP70 resulted largely from its N-terminal domain rather than its C-terminal substrate-binding domain in the context of covalent fusion [22], suggesting that the substrate-interaction forces of chaperones do not necessarily represent the major substrate-stabilizing forces against aggregation.

We previously proposed a *cis*-acting protein-folding helper system, which appears to operate differently from the classical *trans*-acting chaperones [4, 23]. A hallmark feature of the cellular folding environment is that nascent polypeptides are tethered to the cellular macromolecules, such as ribosomes (2000–3200 kDa), membranes, or cotranslationally folded (or prefolded) domains in multi-domain proteins. *De novo* protein folding on these cellular macromolecules has been a major issue in terms of chaperone function [7, 24–26], but the tethering effect of such macromolecules has long been underappreciated. However, based on the robust chaperone-like activity of these macromolecules, as well as a variety of highly soluble proteins, toward various heterologous aggregation-prone proteins in the fusion context (or in *cis*) [27–30], this *cis*-acting chaperone-like type was proposed to play a pivotal role in the folding and aggregation inhibition of endogenous proteins [23]. Consistent with this *cis*-acting model, several lines of evidence indicate that the cytosol-exposed nascent chains tethered to ribosomes are aggregation-resistant and co-translational folding-competent [31–34]. Remarkably, intermolecular repulsive (or destabilizing) forces, such as electrostatic and steric repulsions by the surface charges and excluded volume of cellular macromolecules, were proposed to stabilize their tethered polypeptides against aggregation independently of the attractive intermolecular interactions and their effect on the conformational changes, while the tethered polypeptides can fold based on their own sequence information in the absence of adenosine triphosphate (ATP) consumption [4, 23]. This stabilizing mechanism can underlie the intrinsic chaperone activity of soluble macromolecules. The magnitudes of these intermolecular repulsive forces were suggested to increase corresponding to the size and surface charge of the molecules [22, 23]. Importantly, these two forces have been well known as major factors in stabilizing colloids against aggregation [35, 36]. This stabilizing mechanism well explains the surface-charge effect of HSP90 on anti-aggregation, as well as obvious charge effects on protein solubility [4, 23]. Similarly, entropic bristling and hydration by the excluded volume and charged residues of intrinsically disordered proteins or regions were proposed to solubilize their fused proteins [37, 38]. Moreover, the entropic pulling forces of HSP70 resulting from its excluded volume repulsions were proposed to underlie its diverse functions [39]. Nonetheless, so far, the aggregation inhibition by the intermolecular repulsive forces of cellular macromolecules has been largely ignored; instead, the aggregation inhibition has been explained predominantly in the context of the direct attractive interactions between cellular macromolecules and polypeptides. It should be noted that both action mechanisms act independently and simultaneously.

Importantly, large excluded volume and surface charges are the common intrinsic properties of any type of soluble cellular macromolecule including chaperones. This prompted us to hypothesize that cellular macromolecules exhibit the intrinsic chaperone activity, and thus they act as chaperones for their connected polypeptides irrespective of the connection types between them [4, 40]. To test this hypothesis, we here constructed a *trans*-acting artificial chaperone system. Our results revealed that a model soluble protein exhibits the intrinsic chaperone activity to recapitulate the core features of the classical chaperones, such as aggregation inhibition and folding assistance, upon binding to aggregation-prone proteins in *E. coli*.

## Results

### Design of an artificial chaperone system *in vivo*

To explore the intrinsic chaperone activity of a soluble protein, we designed an artificial chaperone system in which a model soluble protein (named RS-mTEV) specifically binds to a short flanking tag of 7 residues in the substrate proteins (**Fig 1**). As a substrate-binding module of RS-mTEV, we chose a mutant protease domain of tobacco etch virus (mTEV) with no proteolytic activity, but still maintaining the binding affinity for its canonical recognition sequence (ENLYFQG) [41]. This protease domain is marginally soluble when expressed alone at 37 °C [42]. To increase mTEV solubility, it was fused to the C-terminus of *E. coli* lysyl tRNA synthetase (RS; 57 kDa), which is known to be a solubility-enhancing fusion partner [30], resulting in a more soluble RS-mTEV protein (**S1 Fig**). As a client protein of RS-mTEV, enhanced green fluorescent protein (EGFP) was fused to hepatitis B virus X protein (HBx) with intrinsically disordered regions [43], to yield L-EGFP-HBx where “L” denotes the recognition sequence (ENLYFQG). This model system was designed to minimize the direct binding except for the “L” tag between RS-mTEV and its client protein during folding and aggregation processes to assess the intrinsic RS-mTEV chaperone activity. Moreover, the substrate-binding domain (mTEV) is separated as an independent module from the solubility-enhancing module (RS) in RS-mTEV, providing a unique opportunity to distinguish between the contributions of the two modules to RS-mTEV chaperone function later.

**Fig 1.**
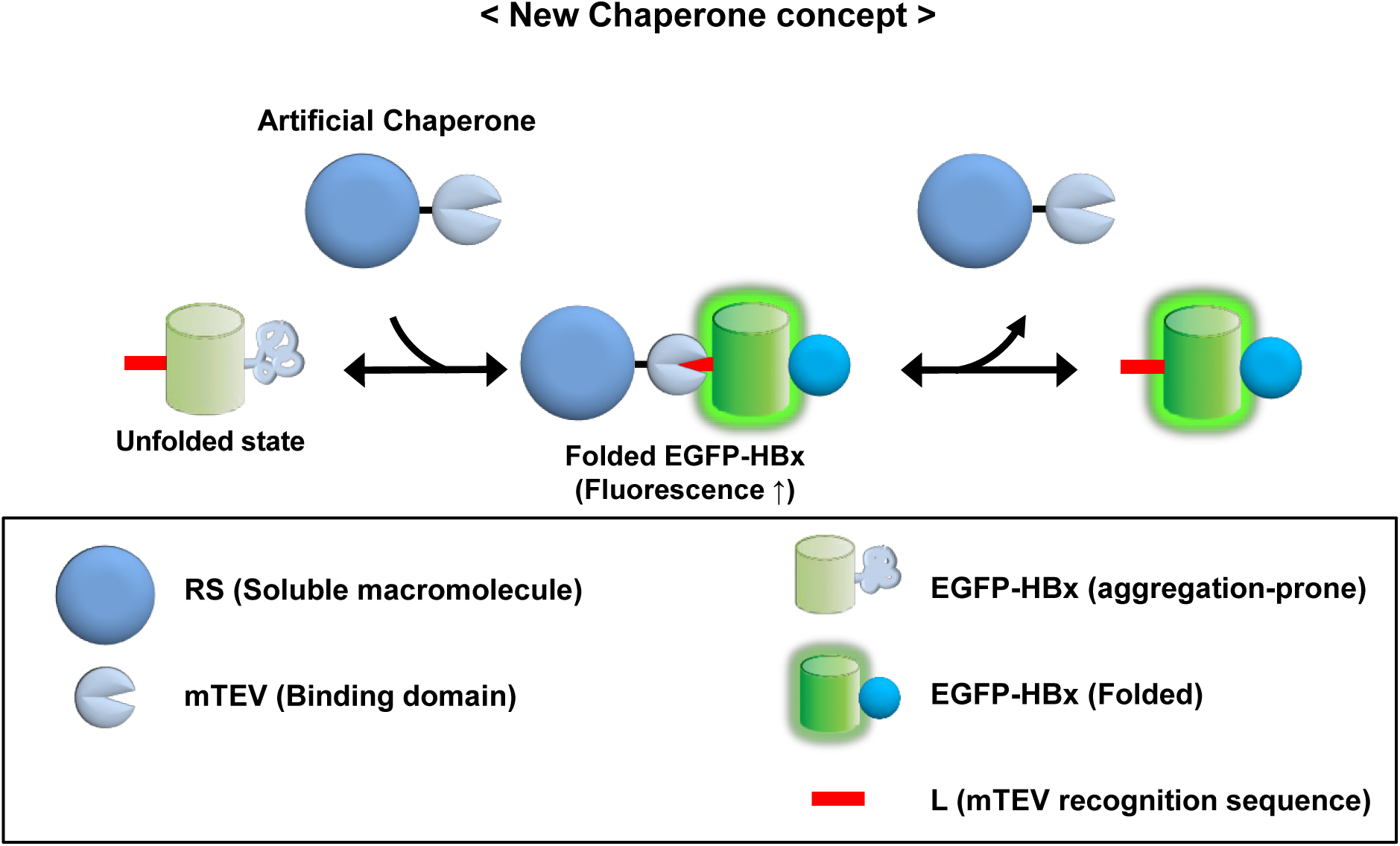
Experimental design for conversion of a soluble protein into a chaperone. Schematic diagram for the construction of an artificial chaperone system to assess the intrinsic chaperone activity of soluble cellular macromolecules. A TEV protease-domain mutant (mTEV) with no proteolytic activity but the binding ability toward its canonical sequence of 7 residues (denoted as “L”; red bar) was fused to the C-terminus of *E. coli* RS, yielding an artificial chaperone, RS-mTEV. EGFP-HBx harboring “L” tag is a client protein of RS-mTEV.

### RS-mTEV acts as a potent chaperone for its client proteins *in vivo*

We investigated the effect of RS-mTEV co-expression on both L-EGFP-HBx solubility and folding in *E. coli* using two co-expression vectors. Information about these vectors is described in more detail (**S2 Fig**). RS-mTEV co-expression markedly increased L-EGFP-HBx solubility by ∼75%, whereas RS co-expression did not increase the solubility (∼16%) similar to the corresponding solubility (∼12%) in background cells containing a mock vector pLysE as a control (**Fig 2a**). We further confirmed that a specific binding of RS-mTEV to the “L” tag in L-EGFP-HBx increased the protein solubility. The residue N171 in mTEV is important for the substrate recognition [41]; therefore, this mutation in RS-mTEV [named RS-mTEV(N171A)] resulted in a significantly impaired substrate-binding ability (**S3 Fig**). Correspondingly, RS-mTEV(N171A) had no detectable solubility-enhancing ability for the substrate protein (**Fig 2a**). Similarly, the solubility of L(m)-EGFP-HBx with the mutation in the “L” tag (ENLYFQG to YNLEFQG) did not respond to RS-mTEV co-expression. Other mutations in the conserved recognition sequence of “L” tag consistently resulted in little or no effect on the protein solubility like L(m)-EGFP-HBx (**S4 Fig**). As expected, EGFP-HBx solubility without the recognition sequence “L” was unaffected by RS-mTEV co-expression (**Fig 2a**). These results clearly demonstrated that RS-mTEV increased the protein solubility via its specific binding to the “L” tag in L-EGFP-HBx *in vivo*. Western blot analysis of the substrate proteins using an anti-GFP antibody was in accordance with their corresponding expression patterns on the above sodium dodecyl sulfate polyacrylamide gel electrophoresis (SDS-PAGE) results. We then investigated the folding quality of the proteins solubilized by RS-mTEV co-expression by measuring EGFP fluorescence intensity in soluble fractions containing the substrate proteins. Solubility enhancement or aggregation inhibition, as important elements for biological relevance, does not necessarily represent proper folding. We observed correlations between EGFP fluorescence intensity and L-EGFP-HBx solubility resulting from RS-mTEV co-expression (**Fig 2a**), indicating that RS-mTEV promoted both solubility and folding of its client protein. The results (**Fig 2a**) revealed that RS-mTEV exhibits the intrinsic chaperone activity.

**Fig 2.**
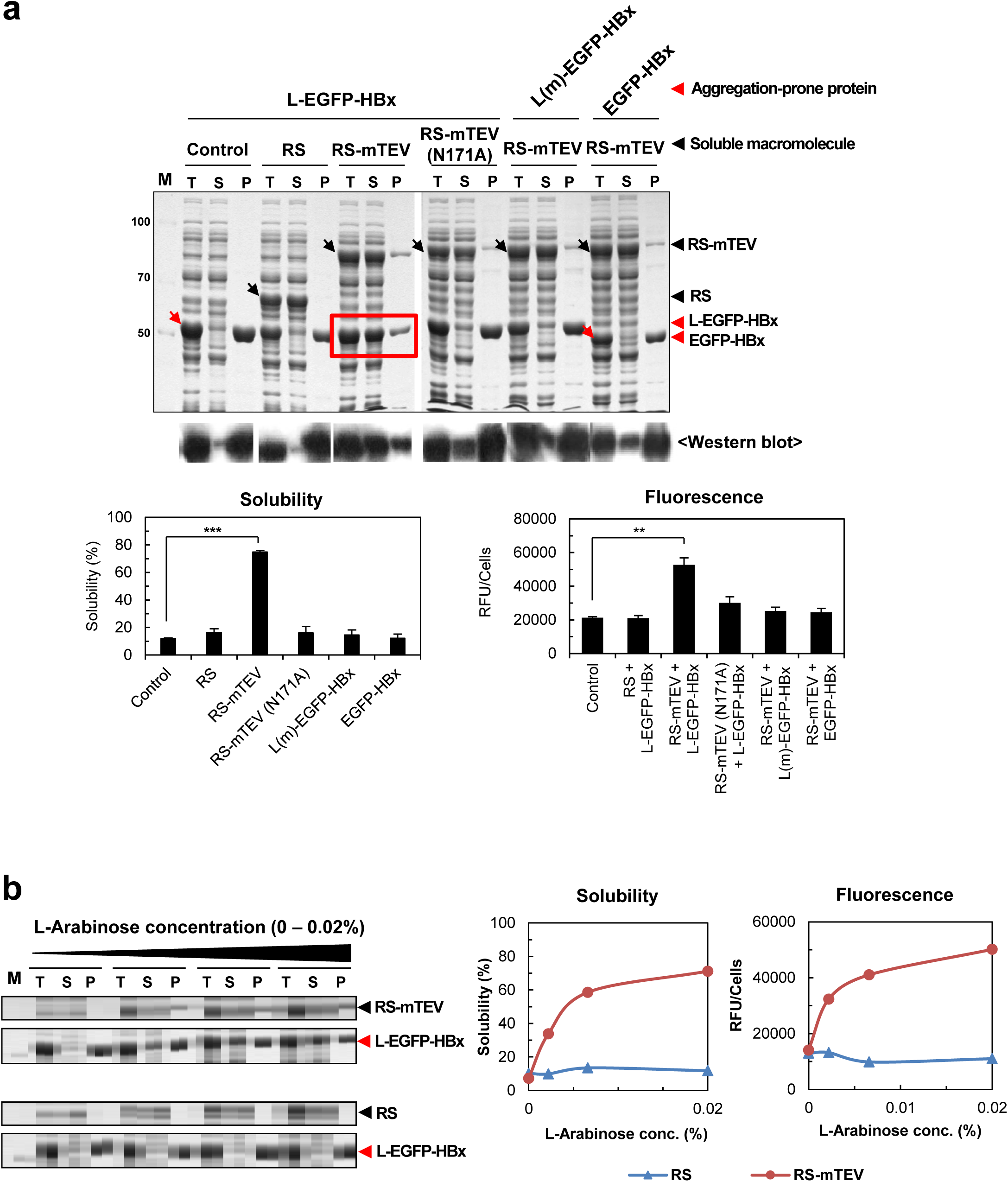
RS-mTEV exhibits potent chaperone activity upon binding to aggregation-prone proteins in *E. coli*. (**a**) Effect of co-expression of RS-mTEV on the solubility and folding of L-EGFP-HBx. RS-mTEV was co-expressed with L-EGFP-HBx, L(m)-EGFP-HBx, and EGFP-HBx, respectively. As negative controls, mock vector pLysE (Control), RS, RS-mTEV(N171A), L(m)-EGFP-HBx, and EGFP-HBx were used. RS-mTEV(N171A) and L(m)-EGFP-HBx harbor the mutations in mTEV and “L” tag, respectively, critical to the substrate protein recognition. Proteins were expressed at 25 °C. The total lysate (T), soluble fraction (S), and pellet (P) of each sample were subjected to SDS-PAGE and western blot analyses. Both solubility on SDS-PAGE and the fluorescence intensity of the EGFP fusion proteins in each soluble fraction were measured and compared. The same analytical methods were used for the following Figs 3-5. Throughout this paper, black and red arrows indicate artificial chaperones (equivalent or control) and substrate proteins, respectively. (**b**) RS-mTEV concentration-dependent chaperone activity. RS-mTEV co-expression was controlled by different concentrations (0–0.02%) of L-arabinose. RS was used as a control.

We further investigated the dosage effects of co-expressed RS-mTEV on L-EGFP-HBx solubility and folding. The increased amounts of co-expressed RS-mTEV protein with increasing L-arabinose concentration (0 %, 0.0022%, 0.0066%, and 0.02%) promoted both L-EGFP-HBx solubility and EGFP fluorescence in an RS-mTEV dosage dependent manner (**Fig 2b**). By contrast, the increase in RS co-expression had no effect on L-EGFP-HBx solubility and EGFP fluorescence under the same condition (**Fig 2b**). These results (**Fig 2**) showed that RS-mTEV was readily converted into a potent chaperone *in vivo* if simply connected to an aggregation-prone protein *in trans*.

### RS-mTEV acts as a chaperone independently of the recognition-tag position

One of the advantages of our artificial chaperone system is that the position of the recognition tag in the substrate proteins can be changed. The intrinsic chaperone activity of RS-mTEV led us to predict that RS-mTEV should act as a chaperone, independently of the position of the recognition tag in the substrate proteins. To test this scenario, the “L” tag was placed either in the middle between EGFP and HBx (EGFP-L-HBx) or at the C-terminus of the protein (EGFP-HBx-L). RS-mTEV co-expression increased the solubility of both EGFP-L-HBx and EGFP-HBx-L (from 49% to 86% and from 12% to 73%, respectively) (**Fig 3**). Consistently, the EGFP fluorescence intensity of the substrate proteins in the soluble fractions was positively correlated with their solubility (**Fig 3**). These results have shown that RS-mTEV acts as a chaperone for its substrate proteins independently of the recognition-tag position, giving further credence to the intrinsic chaperone activity of RS-mTEV. In the cases of L-EGFP-HBx and EGFP-L-HBx, we do not know whether RS-mTEV acted co- or post-translationally. EGFP-HBx-L clearly indicated that RS-mTEV acted at least post-translationally, thereby broadening the generality of our system.

**Fig 3.**
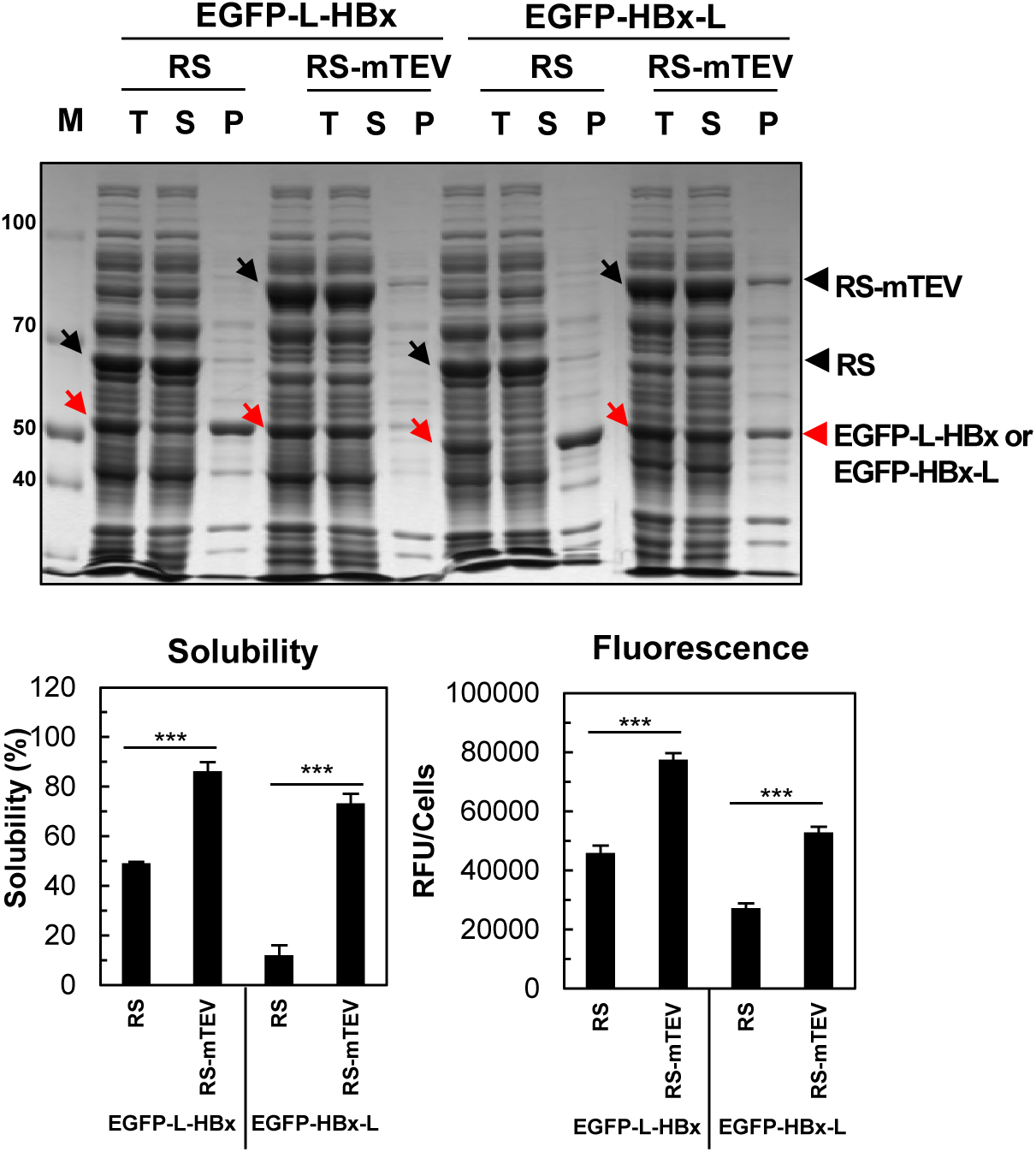
RS-mTEV acts as a chaperone, independently of the recognition tag-position. The “L” tag was placed either in the linker region between EGFP and HBx (EGFP-L-HBx) or at the C-terminus of protein (EGFP-HBx-L). Effect of RS-mTEV coexpression on the solubility and fluorescent intensity of EGFP-L-HBx and EGFP-HBx-L was analyzed as described in Fig. 2. RS was used as a control.

### RS-mTEV is more efficient than the classical chaperones

The representative chaperones, including GroEL-GroES (GroELS), the DnaK-DnaJ-GrpE system (DnaKJE), and TF, have been well known to prevent aggregation and assist protein folding in *E. coli* [7, 9, 26]. Here, we compared RS-mTEV chaperone activity with that of these classical chaperones for L-EGFP-HBx, EGFP-HBx-L, and EGFP-HBx proteins. When co-expressed individually with RS, RS-mTEV, GroELS, DnaKJE, and TF, the corresponding solubility of the 3 substrate proteins were observed to be 16%, 72%, 20%, 81%, and 64% for L-EGFP-HBx, and 12%, 71%, 16%, 82%, and 41% for EGFP-HBx-L, and 10%, 9.8%, 14%, 86%, and 33% for EGFP-HBx (**Fig 4a and c**). The co-expression of the chaperones was confirmed (**S5 Fig**). The results showed that, like RS-mTEV, DnaKJE and TF substantially increased the solubility of L-EGFP-HBx and EGFP-HBx-L to the similar levels, whereas GroELS increased little, similar to RS. Furthermore, the EGFP fluorescence intensities of the soluble extracts were correlated with the corresponding solubility of the target proteins upon co-expression with each chaperone, except for DnaKJE (**Fig 4b**). TF increased both the solubility and folding of client proteins more efficiently than GroELS and DnaKJE. Notably, RS-mTEV was shown to be similar to TF regarding the chaperone activity.

**Fig 4.**
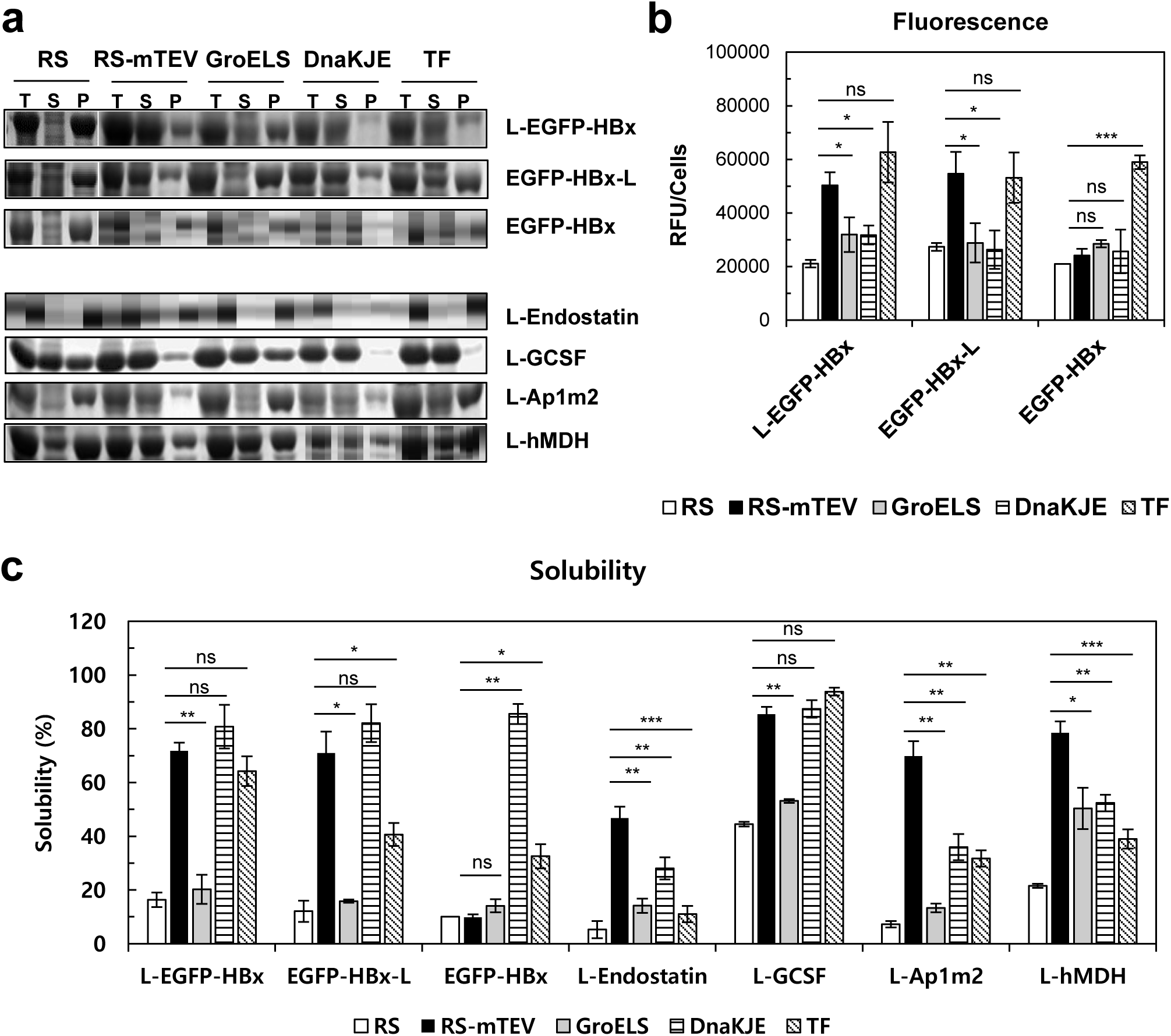
Comparison of RS-mTEV with the representative *E. coli* chaperones. (**a**) Client proteins co-expressed with RS, RS-mTEV, GroELS, DnaKJE, and TF, respectively, were L-EGFP-HBx, EGFP-HBx-L and EGFP-HBx, as well as endostatin, GCSF, Ap1m2, and hMDH. Their expression patterns on SDS-PAGE were highlighted. (**b**) Comparison of EGFP fluorescence of L-EGFP-HBx, EGFP-HBx-L, and EGFP-HBx of the results shown in **a**. (**c**) Comparison of protein solubility of the substrate proteins in **a**.

We additionally compared the solubility-enhancing effects of RS-mTEV with the representative chaperones for different substrate proteins, including human endostatin, granulocyte colony stimulating factor (GCSF), AP-1 complex subunit mu-2 (Ap1m2), and malate dehydrogenase (hMDH) with the “L” tag at their N-termini. These aggregation-prone proteins have been known to be involved in cell proliferation or signaling pathways [44–47]. Upon individual co-expression of RS, RS-mTEV, GroEL/ES, DnaKJE, and TF, the corresponding solubility were 5%, 47%, 14%, 28%, and 11% for endostatin; 44%, 85%, 53%, 87%, and 94% for GCSF; 7%, 70%, 13%, 36%, and 32% for Ap1m2; and 22%, 79%, 50%, 52%, and 39% for hMDH, respectively (**Fig 4a and c**). These results showed that RS-mTEV robustly increased the solubility of all tested proteins, and that its solubility-enhancing activity was higher than or similar to that of the representative chaperones. In contrast to the substrate preferences of the classical chaperones, RS-mTEV provided the chaperone function for all client proteins tested. The overall results (**Fig 4**) indicate that RS-mTEV is more efficient than the classical chaperones.

### RS-mTEV chaperone activity largely results from RS rather than mTEV

As described in Introduction, the substrate-stabilization against aggregation by the surface charges of HSP90, the N-terminal domain of HSP70, and the intermolecular repulsive (or destabilizing) forces of soluble macromolecules appear to act allosterically; long-range chaperone effects exist even in the absence of direct contact with the aggregation-prone regions of the connected polypeptides. This (apparent) allosteric mechanism underlies the concept of the intrinsic chaperone activity of soluble cellular macromolecules. One would therefore expect that RS-mTEV chaperone activity might be mediated by RS after RS-mTEV binding to the “L” tag of client proteins. To test this, we investigated and compared the chaperone effects of three proteins (mTEV without fusion, N-mTEV [N: N-terminal domain (15 kDa) of RS], and RS-mTEV) on L-EGFP-HBx solubility and folding at low-temperature (25 °C), where the solubility of all three proteins is high (**Fig 5a**). Although the three proteins share the same substrate-binding module (mTEV), RS-mTEV was superior to mTEV and N-mTEV at promoting L-EGFP-HBx solubility, whereas N-mTEV chaperone activity was slightly higher than that of mTEV (**Fig 5b and d**). To further confirm RS-mediated chaperone activity, we used a more soluble TEV variant (TEVsw) [48] harboring the same mutation in the TEV domain to block protease activity, yielding mTEVsw. The mTEVsw without fusion was highly soluble, even at 37 °C (**Fig 5a and S1 Fig**). Despite the increased solubility of mTEVsw relative to mTEV, mTEVsw without the RS fusion failed to show detectable chaperone activity for L-EGFP-HBx, whereas RS-mTEVsw consistently increased L-EGFP-HBx solubility (**Fig 5c and d**). The significant amounts of both mTEV and mTEVsw were observed to co-precipitate with L-EGFP-HBx (**Fig 5b and c**). Consistent with the aforementioned allosteric mechanisms, our findings indicate that RS-mTEV chaperone activity results largely from RS rather than mTEV, although mTEV is critical for the substrate binding. Conversely, both the attractive interactions between mTEV module and L-EGFP-HBx and the probable conformational changes of L-EGFP-HBx by the attractive interactions cannot be sufficient to describe the RS-mediated allosteric chaperone effect.

**Fig 5.**
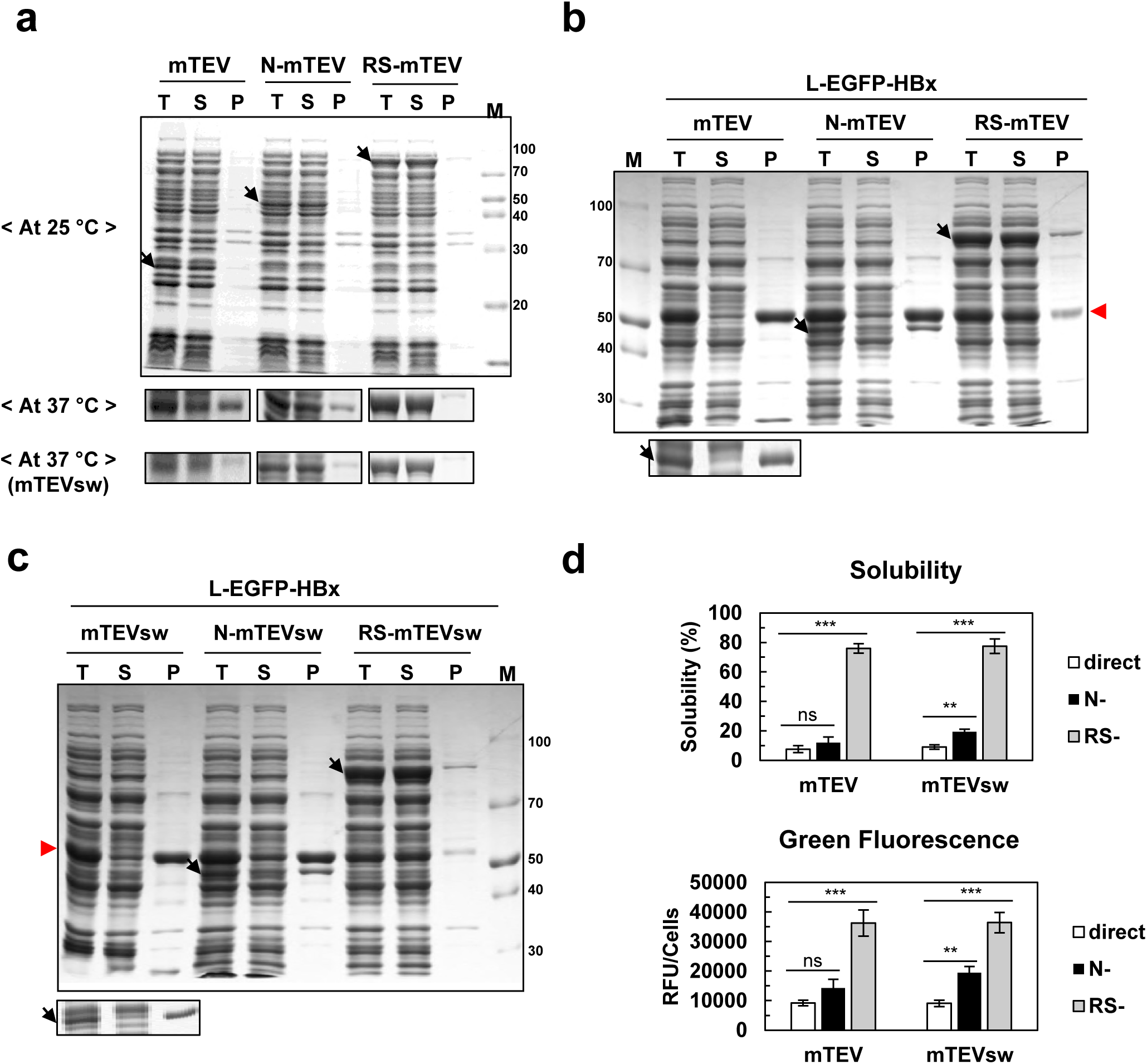
RS-mTEV chaperone activity is largely dependent on RS rather than mTEV. To distinguish between the contributions of RS and mTEV to the RS-mTEV chaperone activity, the chaperone activities of three proteins (mTEV, N-mTEV, and RS-mTEV) and their corresponding more soluble variants (mTEVsw N-mTEVsw, and RS-mTEVsw) were compared. Here N represents the N-terminal domain of RS. (**a**) Solubility of mTEV, N-mTEV, RS-mTEV, mTEVsw, N-mTEVsw, and RS-mTEVsw at 25 °C and 37 °C. (**b**) Comparison of the chaperone activities of mTEV, N-mTEV, and RS-mTEV for L-EGFP-HBx at 25 °C. The mTEV and its fusion variants are indicated by black arrows, and the red arrow indicates L-EGFP-HBx. (**c**) Comparison of the chaperone activity of mTEVsw N-mTEVsw, and RS-mTEVsw at 25 °C under the same conditions as described in **b**. Highlighted bands below main SDS-PAGE data in **b** and **c** represent mTEV and mTEVsw, respectively. (**d**) Solubility and fluorescence intensity of each sample in **b** and **c**.

### *In vitro* refolding experiments support RS-mTEV chaperone function

To characterize RS-mTEV chaperone function more clearly, *in vitro* refolding experiments using EGFP-HBx-L as a substrate protein were performed in the presence and absence of RS-mTEV. GuHCl-denatured EGFP-HBx-L (80 µM) was 50-fold diluted into the refolding buffer containing 2.5 µM RS-mTEV, RS, or phosphate-buffered saline (PBS), and refolding was monitored by following EGFP fluorescence at various time points (0–75 min) at 30 °C. RS-mTEV increased the final refolding yield by ∼ 1. 7 fold high, compared to RS and PBS (**Fig 6a**). Furthermore, consistent with the *in vivo* results (**Fig 2b**), the refolding yields were increased with an RS-mTEV concentration dependence (0–5 µM; **Fig 6b**). By contrast, RS failed to show any detectable chaperone activity, even at the highest concentration tested (**Fig 6a and b**). The final refolding yields of RS-mTEV, RS, and PBS did not converge to the same value (**Fig 6a**), indicating that the difference in the final fluorescence signals resulted from irreversible aggregation of the substrate proteins. This implies that RS-mTEV likely assisted protein folding by preventing off-pathway aggregation. To confirm this possibility, *in vitro* refolding was performed at substrate concentrations 10-fold lowered (**Fig 6c**), where intermolecular aggregation was minimized. Under these conditions, RS-mTEV chaperone activity was substantially attenuated (**Fig 6c**), indicating that RS-mTEV assisted protein folding largely by preventing intermolecular aggregation rather than accelerating the folding rate.

**Fig 6.**
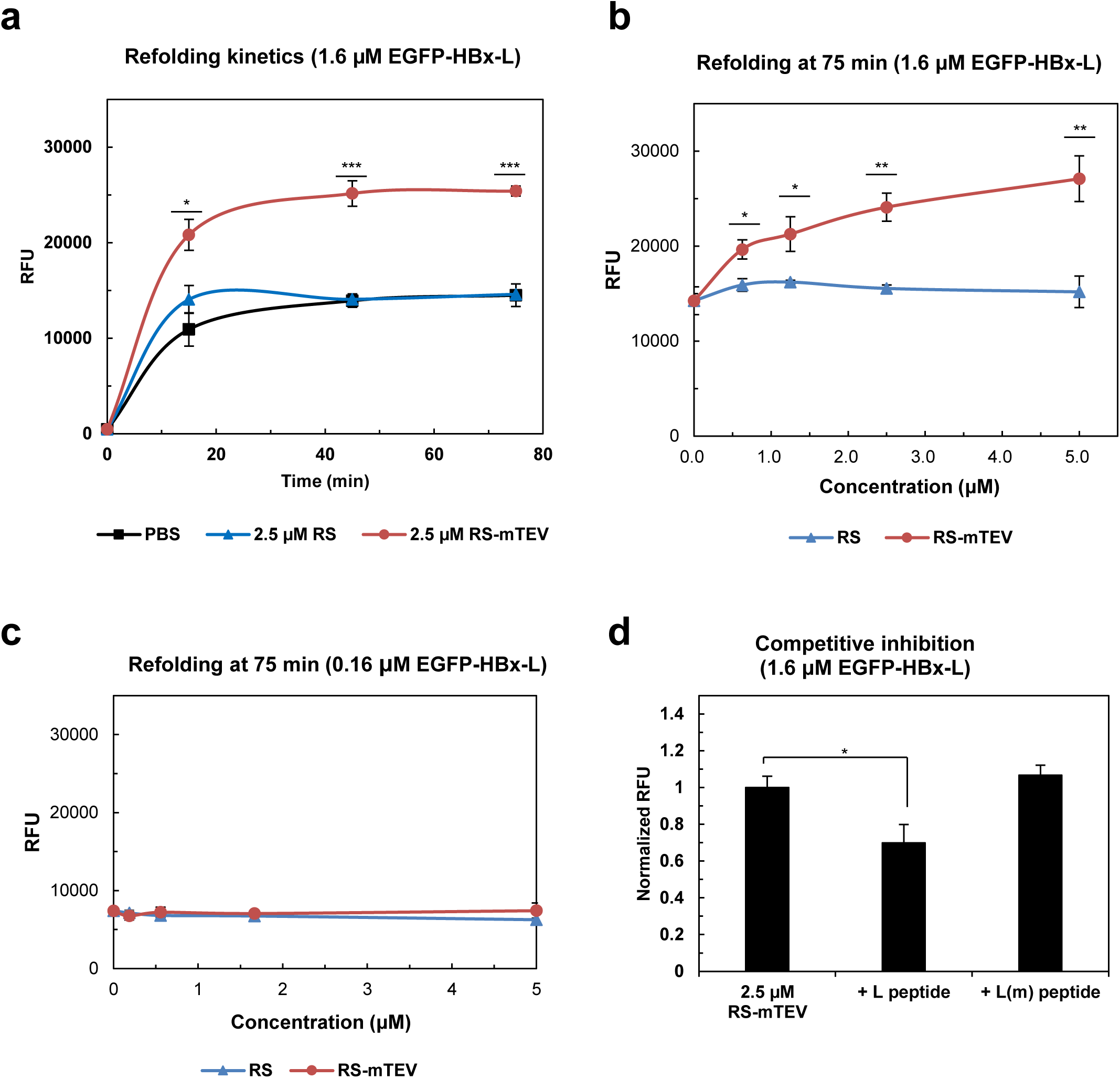
Characterization of the chaperone function of RS-mTEV *in vitro*. (**a**) Refolding kinetics of EGFP-HBx-L (1.6 µM) in the presence of RS-mTEV (2.5 µM) was monitored as a function of time (0, 15, 45, and 75 min). RS and PBS buffer were used as controls. (**b**) Dose-dependent effects of RS-mTEV on EGFP-HBx-L refolding. EGFP fluorescence of the refolded proteins at concentration (0**–**5 µM) of RS-mTEV (or RS) was measured at 75 min after initiation of refolding. (**c**) Loss of RS-mTEV chaperone activity at 10-fold lower substrate concentrations as compared with those in **a**. (**d**) Specific inhibitory effect of the peptide on RS-mTEV chaperone activity. Refolding experiments were similar to those described in **b**, except for the presence of competing (L) or non-competing [L(m)] peptide.

To further confirm *in vitro* RS-mTEV chaperone activity via the specific binding to its canonical recognition sequence in the substrate protein, we investigated the effect of competing peptides on the RS-mTEV chaperone activity. The sequence of the competitive-inhibitor peptide was flanked by TT and GT (TT-ENLYFQS-GT), whereas that of the control peptide represented the inverted form of the canonical recognition sequence (TT-SQFTLNE-GT) harboring a single point mutation in the middle. Addition of the competitive-inhibitor peptide to the refolding buffer abolished RS-mTEV chaperone activity to the level of the control, whereas the control peptide had no inhibitory effect on RS-mTEV chaperone activity (**Fig 6d**). These results demonstrated that *in vitro* RS-mTEV chaperone activity resulted from its specific binding to the canonical recognition sequence.

## Discussion

In this study, we have shown that a soluble protein exhibits the intrinsic chaperone activity in terms of aggregation inhibition and folding assistance. A soluble model protein, RS-mTEV, displayed the robust chaperone activity for its client proteins via a specific binding to the “L” tag of 7 residues (**Figs 2a, 3, and 4**). The fluorescence intensity of EGFP-HBx fusion proteins was followed to assess proper folding because of the absence of an *in vitro* HBx assay [49, 50]. In particular, our artificial chaperone system is suitable to explore the intrinsic chaperone activity of a soluble protein due to the following reasons. The “L” tag is very short and located at the “flanking” regions of the client proteins, minimizing the interactions between RS-mTEV and the client proteins, except for the “L” tag. Moreover, RS-mTEV exhibited a high degree of specificity for the “L” tag (**Figs 2a and 6d**), consistent with a previous report [41]. Similar to such separation of the substrate protein into two parts, RS-mTEV comprises two distinct regions, a solubility-enhancing module (RS) and a client-binding module (mTEV), allowing us to distinguish between the contributions of RS and mTEV to the chaperone activity of RS-mTEV (**Fig 5**). The RS-mediated allosteric chaperone activity in RS-mTEV is in a good accordance with the chaperone activity by the surface charges of HSP90 [21] and the N-terminal domain of HSP70 [22]. Consistently, the intrinsic chaperone activity of soluble macromolecules due to their intermolecular steric and electrostatic repulsions appear to act allosterically; soluble macromolecules can act as chaperones without direct attractive interactions with the aggregation-prone regions of their connected polypeptides [4, 23, 37, 39]. All our findings indicate that RS-mTEV exhibits the intrinsic chaperone activity, which is visible, upon binding to the aggregation-prone proteins.

The concept of the intrinsic chaperone activity of a soluble protein can be generally applicable to cellular macromolecules. This highlights the fundamental importance of our findings. Aggregation-prone polypeptides in the crowded cytosol are physically connected to a variety of cellular macromolecules, including chaperones, through a combination of diverse interactions (covalent/noncovalent, hydrophobic/hydrophilic, transient/permanent, specific/unspecific, direct/indirect, and native/nonnative) [4]. Individual proteins are estimated to continuously interact with the five putative partners in the *E. coli* cytoplasm [51], making quinary interactions with macromolecules inside cells [52]. Our study implies that the above cellular macromolecules potentially act as chaperones for their connected polypeptides irrespective of their connection types. Previously, the intrinsic properties (e.g., excluded volume and surface charges) of soluble proteins and domains were suggested to underlie their robust chaperone activity *in cis* by genetic fusion to aggregation-prone proteins [23]. However, such intrinsic *cis*-acting chaperone activity remains challenging to explore, although it is phenomenologically robust. To facilitate the investigation of such *cis*-acting effects of cellular macromolecules, we initially designed this *trans*-acting system, allowing the independent control of RS-mTEV *in vivo* and *in vitro*. Interestingly, classic chaperones can be converted into potent *cis*-acting solubility enhancers [22, 53, 54], and ribosomes can act as chaperones both in *trans* and in *cis* [27, 34, 55, 56]. Here, RS, a potent solubility enhancer in *cis* [30], provided the *trans-*acting chaperone activity as a component of RS-mTEV (**Fig 5**). All these results show that despite the change in the connection types between the chaperones and their substrates, their chaperone activity persist. Similarly, endoprotease DegP (HtrA) is converted into a chaperone under different conditions [57], with many other protease components previously shown to exhibit chaperone-like activity [58]. Furthermore, various RNAs, highly soluble macromolecules, have been increasingly reported to act as potent chaperones [30, 34, 59–63]. The concept of the intrinsic chaperone activity of the cellular macromolecules in our study can underlie the aforementioned diverse chaperone types. So far, the evolution of the classical chaperones remains largely unknown. Our study implies that soluble macromolecules including protease mutants can be easily converted into chaperones if they have the ability to bind aggregation-prone proteins.

The (apparent) allosteric effect of RS in RS-mTEV on its client protein (**Fig 5**) might be well explained by intermolecular repulsive (or destabilizing) forces exerted by their excluded volume and surface charge. The allosteric modulation of protein aggregation by cellular macromolecules represents a potential mechanism for intervening in aggregation-associated neurodegenerative diseases, as well as for protein solubility *in vivo*. For example, a bulky protein conjugated to amyloid-specific binding dye dramatically inhibits the amyloid formation due to its steric hindrance or excluded volume repulsion [64], consistent with the intrinsic chaperone activity and the allosteric modulation of RS in RS-mTEV. Our findings imply that the cellular macromolecules that bind to the flanking or remote sites away from aggregation-prone regions in proteins and peptides might be potential drug targets for protein aggregation-associated diseases.

Taken together, the present study on the intrinsic chaperone activity of a soluble protein has a huge impact on the field of chaperones, and provides new insights into the generic chaperoning role of cellular macromolecules, which is associated with the cellular protein folding, aggregation inhibition, proteostasis, aggregation-associated diseases, and protein production technology.

## Materials and Methods

### Cloning

We used two different types of vectors for co-expression (pGE and pLysE vectors) (**S2 Fig**). The pGE vectors originated from pGE-LysRS [30], and pLysE vectors were obtained from Merck Chemicals GmbH (Novagen; Darmstadt, Germany). First, the pBAD promoter was inserted into the *Nru*I and *Ava*I restriction sites of pLysE, producing the pLysEpBAD vector with a new *Hpa*I site at the flanking regions of the pBAD promoter. We inserted the sequences for RS, RS-mTEV, variants of RS-mTEV, and the molecular chaperones into the pLysEpBAD vector, respectively. Sequences for client proteins, including EGFP-HBx variants, endostatin, GCSF, Ap1m2, and hMDH, were inserted into the *Nde*I and restriction sites in the multi-cloning site (*Hind*III, *Hind*III, *Sal*I, and *Hind*III, respectively) in the presence or absence of the TEV protease-recognition sequence in the pGE vector. The MSEQ amino acid sequence was inserted at the N-terminus of substrate proteins to increase the expression level of proteins (**S6 Fig**).

### Protein expression

Competent cells [*E. coli* BL21(DE3)] were co-transformed with the aforementioned vectors, cells were grown, and proteins in the cells were analyzed as previously reported[30], with some modifications. The expression of proteins from the pLysEpBAD vector was induced with 0.02% L-arabinose unless otherwise mentioned, followed by culturing at 25 °C for 1.5 h. Then, the cells were treated with IPTG tailored to the expression of each client protein as follows: 50 µM for EGFP-HBx and hMDH, 75 µM for endostatin, 200 µM for GCSF, and 150 µM for Ap1m2. Cells treated with IPTG were cultured at 25 °C for an additional 4 h. The harvested cells were lysed by sonication in PBS, and target-protein solubility in samples was analyzed by SDS-PAGE. All experiments with error bars were performed in triplicate.

### Western blot

Lysate samples were loaded onto polyacrylamide gels and transferred to a polyvinylidene difluoride membrane (ISEQ00010; Millipore, Billerica, MA, USA) according to a previously reported protocol[65]. Anti-GFP (632377; Clontech Laboratories, Mountain View, CA, USA) and anti-Penta His (34660; QIAGEN, Hilden, Germany) were used as primary antibodies, and anti-rabbit IgG (A6154; Sigma-Aldrich, St. Louis, MO, USA) and anti-mouse IgG (A4416; Sigma-Aldrich, St. Louis, MO, USA) were used as secondary antibodies.

### Fluorescence assay of EGFP-fused proteins

The fluorescence of EGFP-HBx variants was measured to investigate the proper folding of proteins. Each soluble fraction of lysed samples was normalized to cellular optical density and added to the well of a 96-well plate (30496; SPL Life Sciences, Gyeonggi-do, Korea). The fluorescent intensity of samples was determined at 485 nm (excitation) and 520 nm (emission), with FLUOstar OPTIMA (BMG Labtech, Cary, NC, USA) used to measure the fluorescence of each well.

### *In vitro* refolding

Purified EGFP-HBx-L proteins in denaturing buffer [50 mM Tris-HCl (pH 7.5), 300 mM NaCl, 6 M GuHCl, 1 mM DTT, and 1 mM EDTA] were supplemented with 50 mM DTT for 30 min before use. The denatured and reduced protein mixtures were 50-fold diluted into refolding buffer [50 mM Tris-HCl (pH 7.5), 150 mM NaCl, and 5 mM MgCl_2_] and incubated at 30 °C. Different concentrations of RS or RS-mTEV were added to refolding buffer along with 1 mM DTT. For each time-course refolding experiment, four individual samples were prepared for monitoring the refolding reaction as a function of time (0, 15, 45, and 75 min) where 0 min corresponds to the sample before dilution of denatured proteins into the refolding buffer. After the initiation of the refolding for 0, 30, 60 min, the samples were centrifuged at 15,000 *x g* for 15 min at 30 °C, making the total refolding time 15, 45, and 75 min, respectively. Green fluorescence intensity in the supernatant of each sample was measured. We used RS-mTEVsw for the refolding experiments, which exhibits better stability and solubility than its prototype [48]. When testing 10-fold lower concentrations of EGFP-HBx-L, 1 mg/mL bovine serum albumin was added to the refolding buffer to reduce loss of the substrate protein due to nonspecific adsorption. All refolding experiments were performed in triplicate.

### Competitive inhibition of refolding

Peptides (5 mM; TT-ENLYFQS-GT and TT-SQFTLNE-GT) were dissolved in 100% dimethyl sulfoxide and used to inhibit the chaperone effect of RS-mTEV in *in vitro* refolding assays.

### Electrophoretic mobility shift assay (EMSA)

Serially diluted and purified RS-mTEV, RS-mTEVsw, and RS-mTEV(N171A) were mixed with 2.5 µM of purified EGFP-L. They were then incubated at room temperature for 30 min and separated on Native-PAGE gel. Before staining with Coomassie brilliant blue (EBP-1011; Elpis Biotech, Daejeon, Korea), a fluorescence image of the gel was captured.

### Statistical analysis

The error bars in each graph represent the mean ± standard deviation of results obtained from triplicate experiments. Statistical significance was analyzed using Student’s *t* test. A two-tailed P-value was considered statistically significant at P < 0.05.

## Data availability

All data generated or analyzed during this study are included in this published article (and its supplementary information files).

## Acknowledgments

We thank Helena Berglund for kindly providing the plasmid encoding an engineered Tobacco Etch Virus protease domain.

## Abbreviations

Ap1m2: AP-1 complex subunit mu-2
DnaKJE: DnaK-DnaJ-GrpE
EGFP: Enhanced Green Fluorescent Protein
GCSF: Granulocyte colony stimulating factor
GroELS: GroEL-GroES
HBx: Hepatitis B virus X protein
hMDH: Malate Dehydrogenase from *Homo sapiens*
HSP: Heat Shock Protein
IPTG: isopropyl β-D-1-thiogalactopyranoside
mTEV: Tobacco etch virus protease domain with C151A mutation
mTEVsw: Tobacco etch virus protease domain variant with C151A mutation
RS: Lysyl-tRNA synthetase of *Escherichia coli*
SDS-PAGE: Sodium dodecyl sulfate polyacrylamide gel electrophoresis
TF: Trigger Factor

## Author contributions

Conceptualization: Seong Il Choi

Formal analysis: Soon Bin Kwon, Hotcherl Jeong, Kyun-Hwan Kim, Baik L. Seong, Seong Il Choi

Funding acquisition: Baik L. Seong

Investigation: Soon Bin Kwon, Kisun Ryu

Methodology: Soon Bin Kwon, Kisun Ryu, Keo-Heun Lim, Seong Il Choi

Supervision: Baik Lin Seong, Seong Il Choi

Validation: Soon Bin Kwon

Visualization: Soon Bin Kwon, Ahyun Son, Seong Il Choi

Writing-original draft: Soon Bin Kwon, Ahyun Son, Baik L. Seong, Seong Il Choi

## Supporting information

**S1 Fig.**
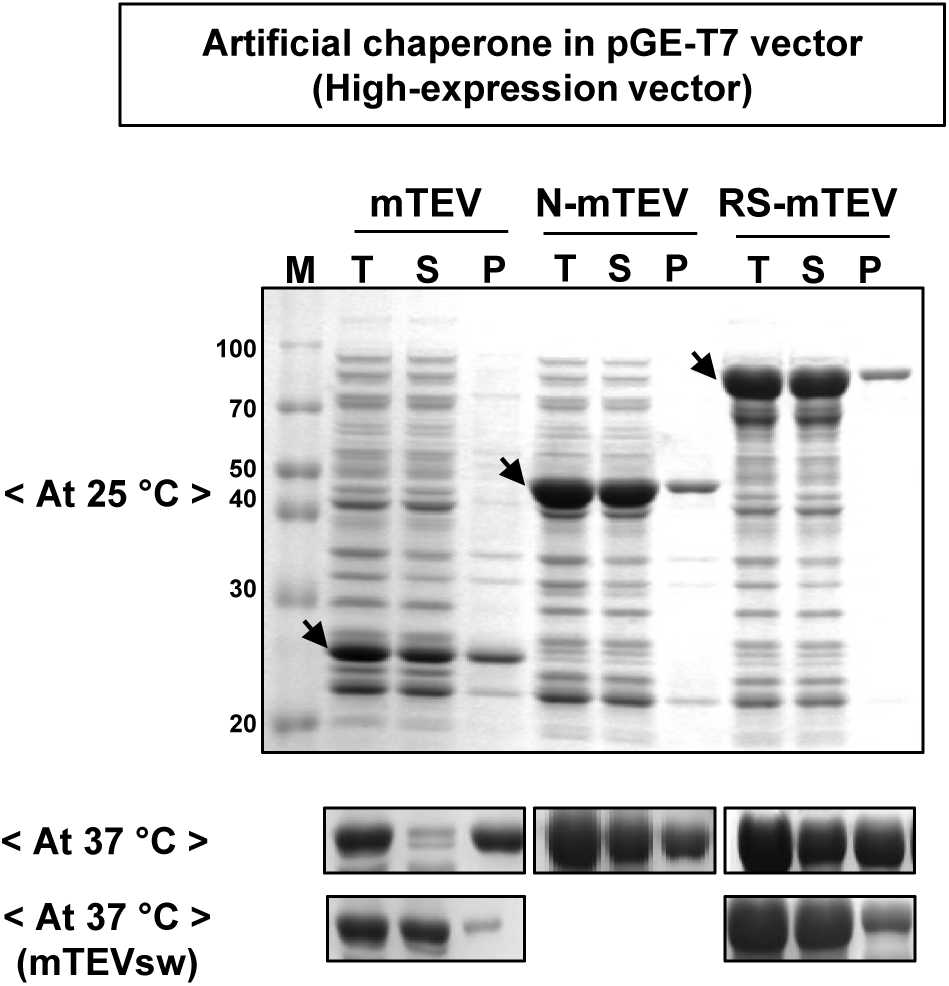
Expression of mTEV and its derivatives in *E. coli.* mTEV, N-mTEV, and RS-mTEV were expressed in *E. coli* at 25 °C and 37 °C. N and RS represent the N-terminal domain of *E. coli* LysRS and the whole LysRS, respectively. More soluble mTEV variant (mTEVsw) and its derivative (RS-mTEVsw) were also expressed in *E. coli* at 37 °C. The solubilities of the three mTEV variants expressed at 25 °C were similar, whereas that of mTEV decreased at 37 °C. mTEVsw alone was highly soluble even at 37 °C.

**S2 Fig.**
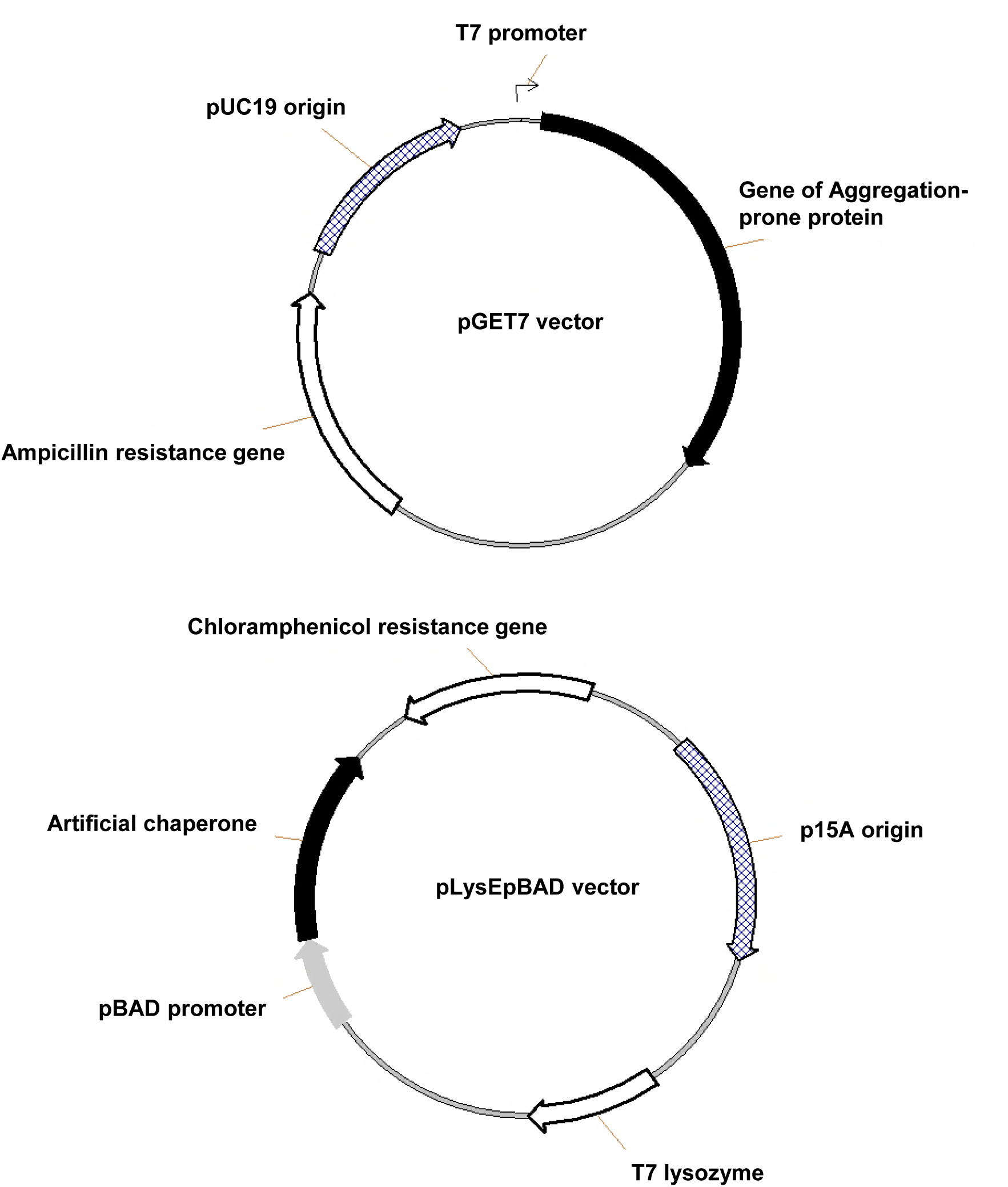
Diagram of co-expression vectors used for aggregation-prone proteins and artificial chaperones. pGET7 vector used for the expression of aggregation-prone substrate proteins harbors an ampicillin-resistance gene and a pUC19 origin of replication. Protein expression under control of T7 promoter was induced by IPTG. pLysEpBAD used for chaperone expression carries a chloramphenicol-resistance gene and a p15A origin of replication. Expression of artificial chaperones under control the pBAD promoter was triggered by L-arabinose.

**S3 Fig.**
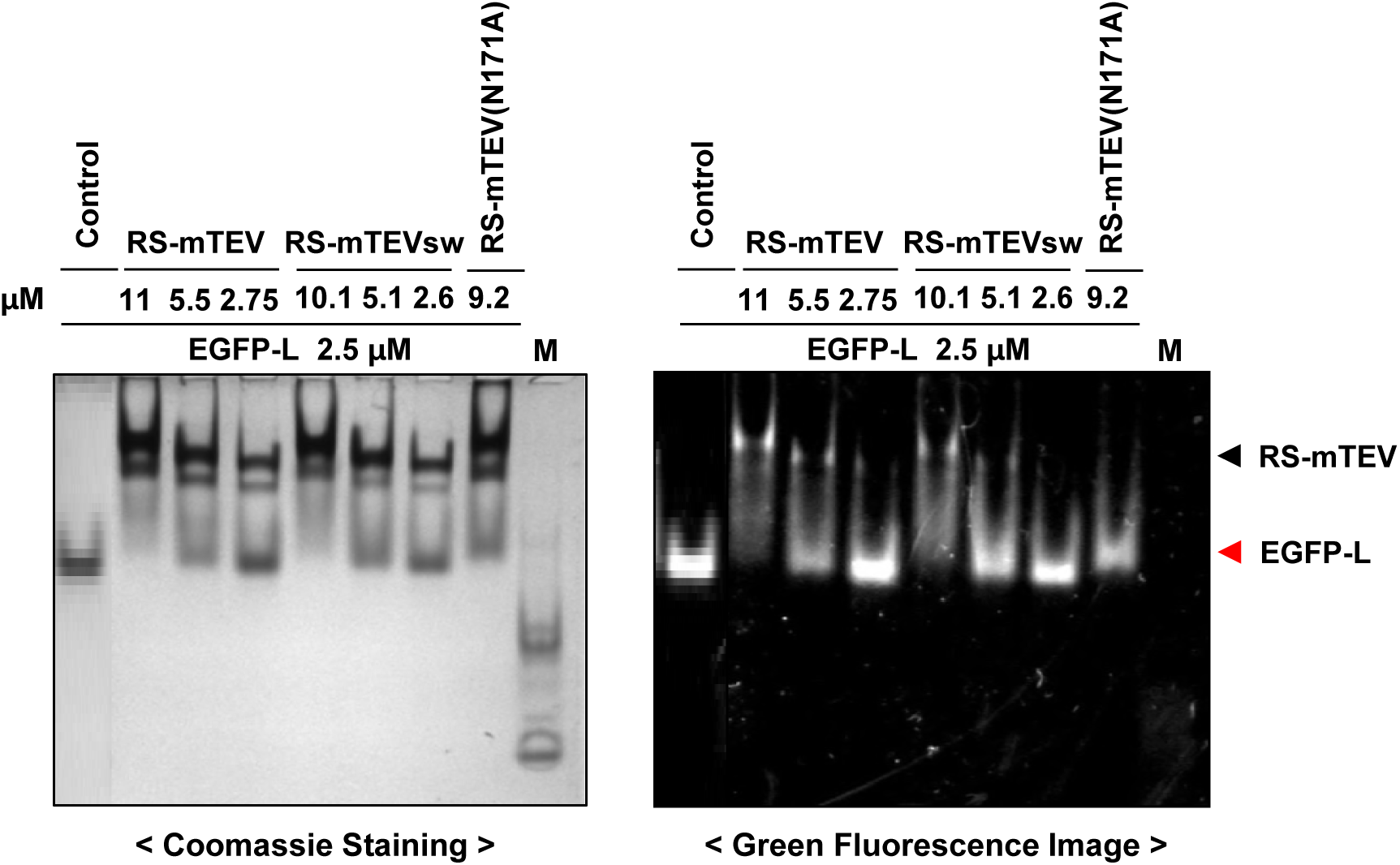
Interaction of the “L” tag with RS-mTEV, RS-mTEVsw, and RS-mTEV (N171A). EGFP-L was mixed with varying concentrations of RS-mTEV, RS-mTEVsw, or RS-mTEV (N171A), and then the binding between them was analyzed by mobility shift on native PAGE. Fluorescence images were first obtained (right), and then staining with Coomassie brilliant blue (left) was performed.

**S4 Fig.**
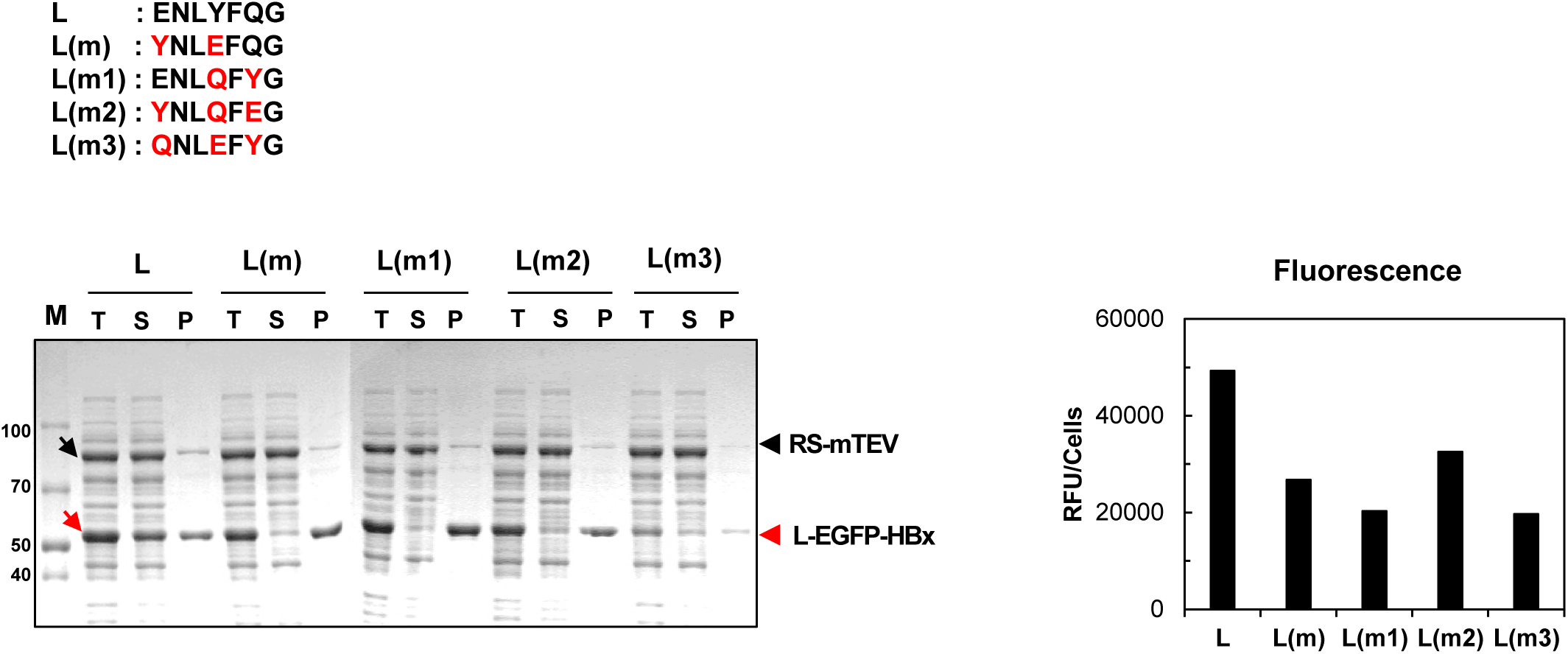
Mutation of the “L” tag in L-EGFP-HBx alters interaction with RS-mTEV. A mutation was introduced in the conserved residues of the “L” tag, and the resulting mutant variants [L(m), L(m1), L(m2), and L(m3)] were attached to the N-terminus of EGFP-HBx, respectively. These proteins were co-expressed with RS-mTEV in *E*. *coli*. Mutated sequences in the “L” tag are indicated in red. Expressed proteins were analyzed by SDS-PAGE and verified by fluorescence (histogram, right).

**S5 Fig.**
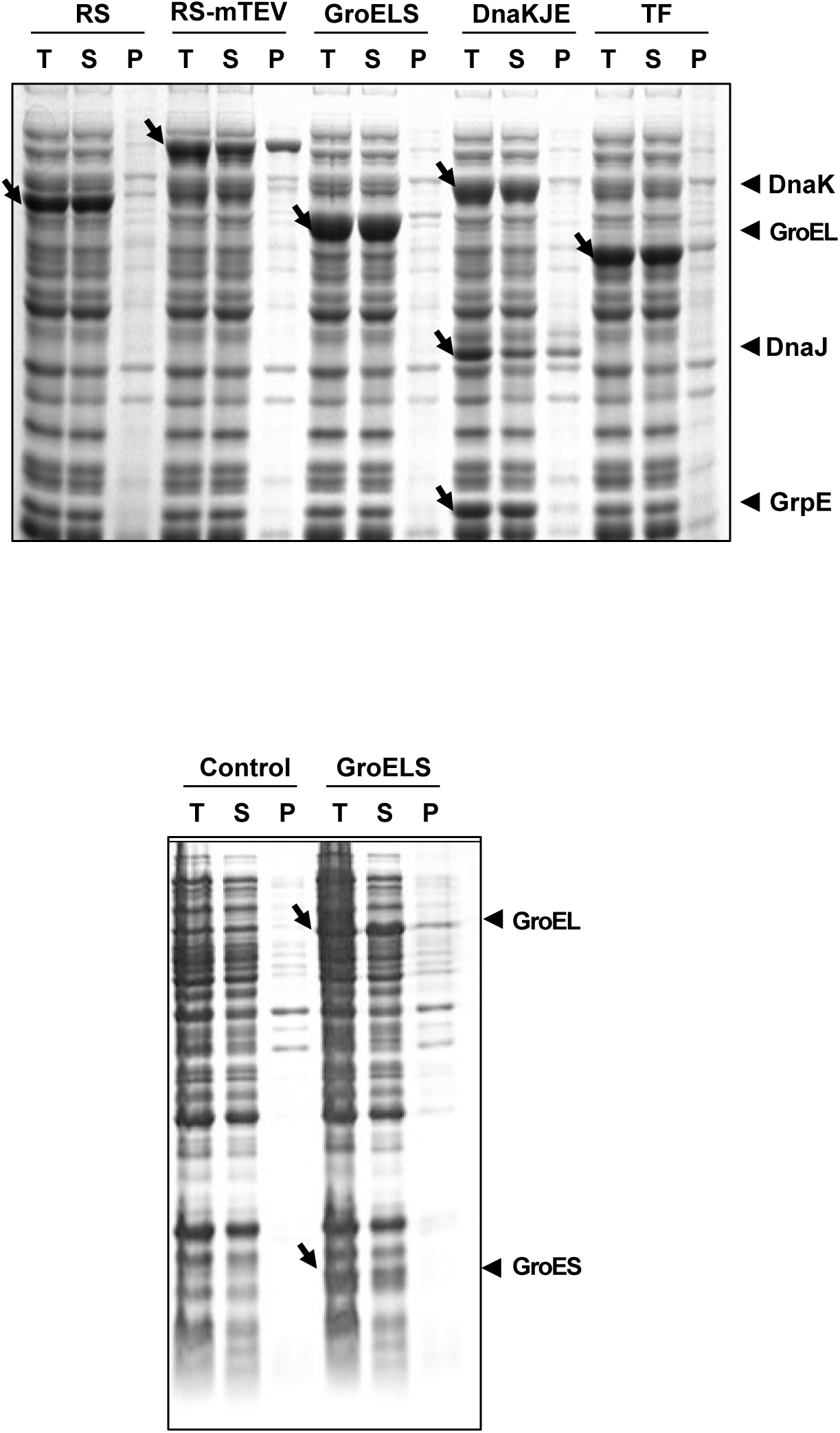
Confirmation of the expression of RS, RS-mTEV, and molecular chaperones. RS, RS-mTEV, GroELS, DnaKJE, and TF were expressed in *E. coli*, and their expression was analyzed SDS-PAGE. Each target band is indicated by an arrow. In the case of GroELS, the SDS-PAGE (down) was added to clearly see the expression of groES, a relatively small sized protein, which was run through in the upper SDS-PAGE obtained after a long running time for a good resolution.

**S6 Fig.**
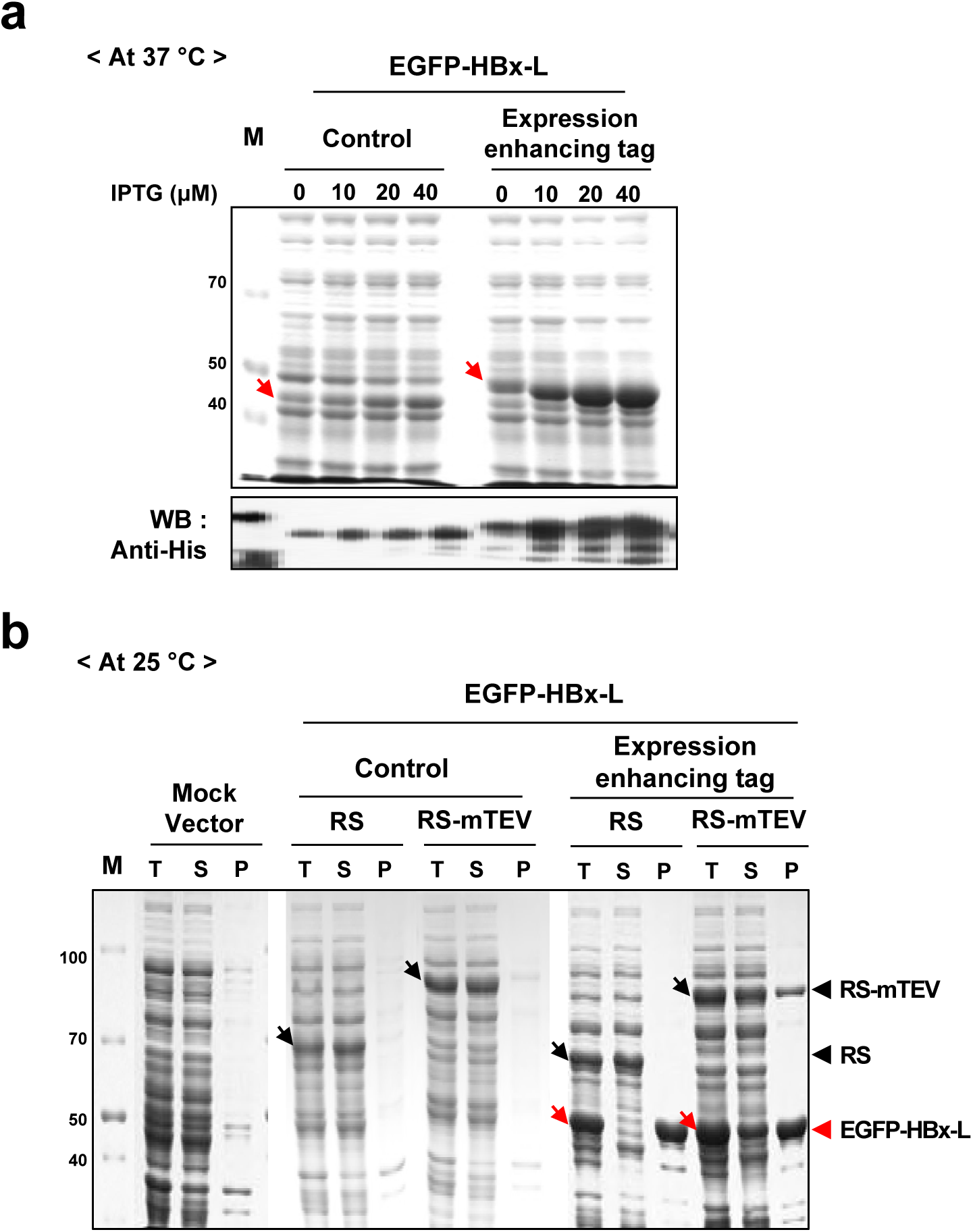
The N-terminal MSEQ tag increases the expression of the substrate proteins of RS-mTEV. (**a**) EGFP-HBx-L in the presence or absence of this tag was expressed in *E. coli* at 37 °C and induced at various IPTG concentrations (0–40 µM). Total lysates of each sample were analyzed by SDS-PAGE and western blot. Red arrows indicate EGFP-HBx-L expression. (**b**) EGFP-HBx-L in the presence or absence of the tag was expressed in *E. coli* at 25 °C (induced by 100 µM IPTG) along with RS or RS-mTEV co-expression, followed by SDS-PAGE analysis.

